# Microglial morphological complexity in the piriform cortex is associated with olfactory aversion following chronic stress

**DOI:** 10.1101/2025.08.14.670114

**Authors:** Kai Clane Belonio, Zach Fyke, Eyerusalem S. Haile, Tahniat Nadeem, Joseph D. Zak

## Abstract

Olfactory anhedonia and heightened aversion to unpleasant odors are well-documented features of depression in humans, yet the neural mechanisms linking chronic stress to altered olfactory perception remain poorly understood. We used the Unpredictable Chronic Mild Stress (UCMS) paradigm to examine how chronic stress affects olfactory avoidance behavior and glial cell morphology across multiple olfactory brain regions in male and female mice. UCMS-treated mice showed increased avoidance of aversive odorants in an odorized light/dark box assay, consistent with heightened aversive reactivity to odors following chronic stress. Using immunohistochemistry, we assessed microglial morphology and astrocyte density across six olfactory and limbic brain regions. Chronic stress produced region-specific glial remodeling: astrocyte counts were selectively elevated in the medial amygdala, and microglial process complexity was increased in the anterior olfactory nucleus and anterior piriform cortex. Microglial morphological complexity in the anterior piriform cortex was correlated with individual odor avoidance scores. These findings reveal that chronic stress induces regionally specific glial plasticity within olfactory sensory and affective networks and suggest that microglial remodeling in piriform cortex may contribute to stress-related changes in olfactory perception.

**Significance Statement:** Changes in sensory perception frequently accompany depression. While previous studies have implicated neuroinflammation in depression-related dysfunction within cortical and limbic structures, little is known about how chronic stress affects glial cells in olfactory processing regions. Here, we show that chronic stress induces glial remodeling in key olfactory areas, including the olfactory bulb, anterior piriform cortex, and medial amygdala, and that these changes correlate with heightened behavioral avoidance of aversive odors. These findings suggest that glial plasticity in sensory networks contributes to affective alterations in olfactory perception, revealing a potential mechanism by which mood disorders can influence sensory experience. This work advances our understanding of the neuroimmune basis of sensory-affective integration.

## Introduction

Chronic stress is a significant risk factor for the development of depressive disorders, and its effects on brain function extend well beyond classic limbic and cortical targets (Agid et al., 2000; Baumeister et al., 2014). Depression affects approximately 5.7% of the world population and is associated with perceptual and cognitive symptoms, including reduced concentration, anhedonia, and changes in appetite, that reflect broad disruption of sensory and affective processing (WHO, 2025; Wu et al., 2025). Among these, alterations in olfactory perception are increasingly recognized as a clinically relevant feature of depressive disorders. Depressed patients exhibit reduced affinity for pleasant odorants, heightened aversion to unpleasant odors, impaired odorant discrimination, and structural changes in the olfactory bulb (Atanasova et al., 2010; Athanassi et al., 2021; Kamath et al., 2018; Naudin et al., 2012; Negoias et al., 2010; Pabel et al., 2018). These findings suggest that chronic stress disrupts olfactory processing at the cellular level, yet the mechanisms underlying this disruption remain poorly understood.

One candidate mechanism is stress-induced glial remodeling. Chronic stress activates both astrocytes and microglia, alters neurogenesis, and elevates inflammatory markers across multiple brain regions (Blossom et al., 2020; Du Preez et al., 2021; Li et al., 2015; Rahimian et al., 2024). In regions such as the hippocampus and prefrontal cortex, this neuroimmune response has been mechanistically linked to behavioral correlates of depression: microglial morphological changes occur rapidly following stress exposure (Kreisel et al., 2014), microglial activation can be broadly induced by chronic stress across multiple brain regions (Calcia et al., 2016), and pharmacological targeting of microglial activation rescues stress-induced behavioral deficits (Farooq et al., 2012; Guo et al., 2022; Tabassum et al., 2022). Astrocytes and microglia do not act independently in this context, as activated microglia can induce neurotoxic reactive states in astrocytes, suggesting that the two cell types may be co-regulated under conditions of chronic stress (Liddelow et al., 2017; Yirmiya et al., 2015). Whether similar glial changes occur in olfactory processing regions and whether they contribute to stress-related alterations in odor-driven behavior have not been examined.

This gap is notable given that glial cells play active roles in normal olfactory function. Astrocytes in the olfactory bulb, in the glomerular and external plexiform layers, regulate odor-evoked excitability through K⁺ clearance and CB₁ receptor-dependent endocannabinoid signaling, shaping the output of mitral and tufted cells (Bellot-Saez et al., 2017; De Saint Jan and Westbrook, 2005; Gómez-Sotres et al., 2024). In downstream regions, astrocytes modulate the integration of chemosensory information by clearing neurotransmitters and mediating gliotransmitter-dependent synaptic regulation, thereby influencing plasticity within networks that guide odor-driven behavior (Martin-Fernandez et al., 2017; Noriega-Prieto et al., 2014; Yang et al., 2025). Microglia, meanwhile, regulate adult neurogenesis in the olfactory bulb through phagocytosis of newborn neurons and dendrites, and their CX3CR1-dependent signaling shapes the integration of adult-born neurons into olfactory processing (Lazarini et al., 2012; Denizet et al., 2016; Reshef et al., 2017; Wallace et al., 2020; Ung et al., 2021; Duan et al., 2020). Stress-induced remodeling of either cell type could therefore alter sensory processing at multiple levels of the olfactory hierarchy. Consistent with this, chronic mild stress in rodents reduces olfactory neurogenesis and alters anxiety-related behavior, suggesting that olfactory regions are sensitive to the effects of chronic stress even in the absence of direct neuroinflammatory challenge (Mineur et al., 2006; Mineur et al., 2007).

We therefore used the Unpredictable Chronic Mild Stress (UCMS) model to investigate how chronic stress affects behavioral responses to aversive odorants and to characterize changes in astrocyte density and microglial morphology across six olfactory and limbic brain regions. By linking regional glial changes to individual differences in odor avoidance behavior, we aimed to identify candidate cellular loci through which stress-induced neuroimmune remodeling may contribute to altered olfactory perception.

## Materials and Methods

### Animals

Twenty-three adult male and female C57Bl6/J mice were used in this study and were randomly sorted into the control group (*n* = 11) or the stressed group (*n* = 12). Both groups were housed in groups of 3-4 in standard laboratory cages (19 cm x 30 cm x 13 cm) at the University of Illinois Chicago Biological Research Laboratory. All mice experienced a 12-hour light/dark cycle at a temperature of 22 ± 2 °C and had access to food and water *ad libitum* except during the Restraint stressor (**Table 1**).

**Table 1.** UCMS protocol stressors.

| Stressor | Description |
| --- | --- |
| Social stress | Mice are placed into a cage that has been previously occupied by another group of mice of the opposite sex for at least two days. |
| Wet cage | A cage without bedding or enrichment was filled 1 cm deep with water. |
| Dampened bedding | 200 mL of water was added for every 100 g of bedding. |
| Empty cage | Cage with no bedding, or enrichment. |
| Tilted cage | Cage was tilted at 45°. |
| Restraint | Mice are placed in a restraining device that prevents movement but still allows access for respiration and air circulation. |

### Ethics Oversight

All experiments were performed in accordance with the guidelines set by the National Institutes of Health and approved by the Institutional Animal Care and Use Committee at the University of Illinois Chicago (protocol 24-167).

### Unpredictable Chronic Mild Stress (UCMS)

Stressed mice were subjected to a series of different mild stressors five days a week for four consecutive weeks. Stressed animals were subjected to one mild stressor per day for four consecutive hours. All stress procedures were performed during the light cycle in a different room from the housing room. Detailed descriptions of the stressors used are in **Table 1**. The order of the stressors was randomized to maintain unpredictability. The experimental schedule for the UCMS model is outlined in **Table 2**. For stressors involving water, the water was kept at a stable, warm temperature (30 °C). Afterwards, mice were placed into a heated recovery cage and returned to their home cage after rejuvenation.

**Table 2.** UCMS protocol schedule.

|  | Day 1 | Day 2 | Day 3 | Day 4 | Day 5 |
| --- | --- | --- | --- | --- | --- |
| Week 1 | Social stress | Wet cage | Dampened sawdust | Empty cage | Wet cage |
| Week 2 | Tilted cage | Empty cage | Restraint | Dampened sawdust | Tilted cage |
| Week 3 | Social stress | Wet cage | Restraint | Empty cage | Dampened sawdust |
| Week 4 | Tilted cage | Wet cage | Empty cage | Social stress | Restraint |

### Behavioral Assessments

Both the control and stressed groups of mice experienced each behavioral assessment. Testing occurred during the light phase, between 8 AM and 2 PM, in a room separate from the housing room. After habituation to each assessment arena, testing occurred one week before and one week after the stressed group experienced UCMS, as outlined in **Figure 1** and **Table 3**.

**Figure 1.**
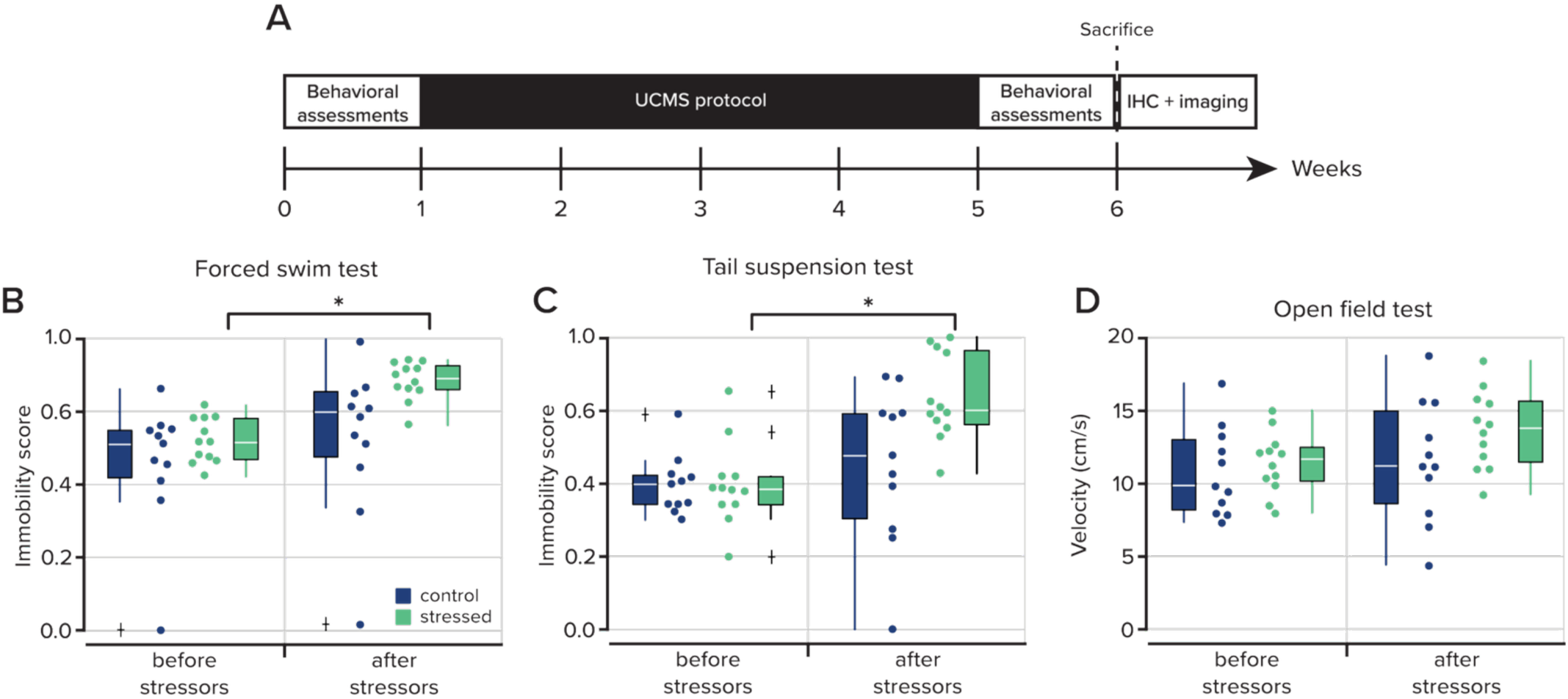
Stress induces a depressive-like state in mice. **A.** Experimental timeline of the UCMS protocol and the behavioral assessments. After the behavioral assessments following UCMS, mice were sacrificed for immunohistochemistry. **B.** The immobility score (see *Methods*) in the Forced Swim Test for the control (*n* = 11) and stressed (*n* = 12) groups from before and after exposure to the UCMS protocol stressors (*p* = 4.883 x 10^-4^, signed-rank test). **C.** The immobility score in the Tail Suspension Test before and after exposure to stressors (*p* = 4.883 x 10^-4^, signed-rank test). **D.** Open Field Test results plotted as the mean velocity of experimental groups before and after exposure to UCMS. Dots represent experimental replicates. The white line in the box represents the median, the box edges indicate the 25th and 75th percentiles, respectively. The whiskers extend to the most extreme data points. Outliers are plotted individually using the ‘+’ marker symbol.

**Table 3.**
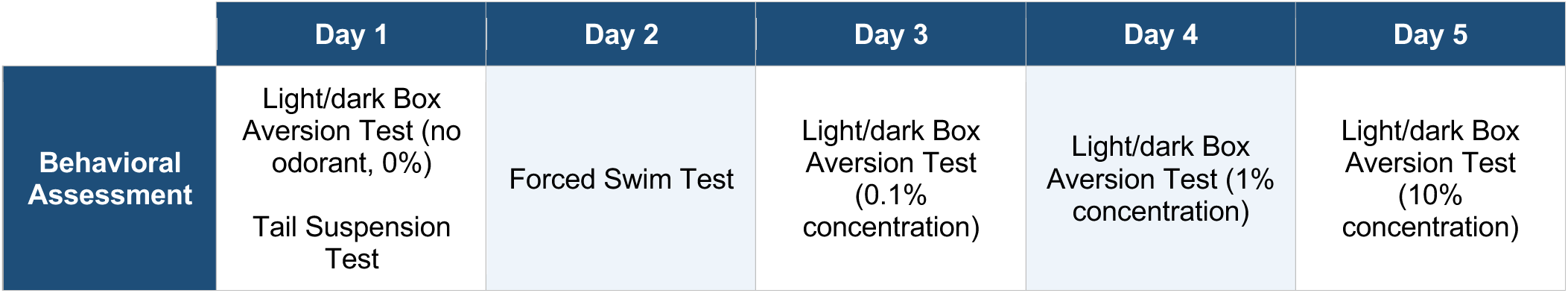
Sample behavioral schedule. These tests were performed one week apart: once before and once after implementing the UCMS protocol. The odorant used was either Trimethylamine or Isopentylamine (see *Light/dark Box Odor Aversion Test* for details).

|  | Day 1 | Day 2 | Day 3 | Day 4 | Day 5 |
| --- | --- | --- | --- | --- | --- |
| Behavioral Assessment | Light/dark Box Aversion Test (no odorant, 0%)<br>Tail Suspension Test | Forced Swim Test | Light/dark Box Aversion Test (0.1% concentration) | Light/dark Box Aversion Test (1% concentration) | Light/dark Box Aversion Test (10% concentration) |

### Forced Swim Test

A clear glass cylinder (30 cm x 15 cm x 15 cm) was filled with room-temperature (23-25 °C) water to a height of 15 cm. Mice were placed at a random point in the water and left to swim for 6 minutes. Videos of each session were manually scored by noting the time the mouse spent immobile in the water. The first two minutes of the session were habituation, and only the last four minutes were scored. Raw scores in seconds were normalized using the formula 1-((value - minimum) / range) and displayed as the Immobility Score. Statistical comparisons were computed in MATLAB using the rank-sum test for unpaired samples and the signed-rank test for paired samples. Data are given as mean ± standard error of the mean (SEM).

### Tail Suspension Test

Mice were suspended at a height of about 20-25 cm using lab labeling tape 3-4 cm from the base of the tail. Plastic cylinders were placed around the mouse’s tail to prevent tail climbing behavior. Videos of each six-minute session were manually scored by noting the time the mouse spent motionless in the air. The full six minutes of the session were scored. Raw scores in seconds were normalized using 1-((value - minimum) / range) and displayed as the Immobility Score. Statistical comparisons were done using the rank-sum test for unpaired samples and the signed-rank test for paired samples. Data are given as mean ± SEM.

### Open Field Test

An opaque 51 cm x 56 cm x 16 cm arena was used for behavioral assessment, and a clear acrylic board was placed over the arena to prevent escape and allow for video recording. Mice were consistently placed in the same corner of the arena and allowed to freely roam for 10 minutes. The arena was cleaned with 70% ethanol between animals. Behavior was recorded using the software IC Capture at 10 frames per second using a USB camera (DFK 42BUC03, The Imaging Source) with an accessory camera lens (IP/CCTV 13VM308ASIRII, Tamron). Videos were further processed in ImageJ (Schindelin et al., 2012) and with custom MATLAB scripts that measured the mouse’s location from the midpoint of its body and tracked its location and average velocity within the arena. Statistical comparisons were done using the rank-sum test for unpaired samples and the signed-rank test for paired samples. Data are displayed as mean ± SEM.

### Odorized Light/Dark Box Test

The behavior apparatus consisted of two 18 cm x 30 cm x 12 cm chambers connected by a 6 cm diameter tube. The light chamber was semi-transparent, while the dark chamber was painted black with acrylic paint. The dark chamber had two air inlets that allowed both air and odor to flow into the chamber at a constant rate of <100 mL/min. The apparatus was placed in a larger sound- and lightproof chamber lit by LED lights. During testing, an opaque board was placed on top of the dark chamber to further dim its interior, and a clear acrylic board was placed over the entire apparatus to keep both mice and odor within the chambers and to allow for video recording. Mice were initially placed into the dark chamber and allowed to explore the apparatus for five minutes, at which point the air inlet in the dark chamber was switched to the odor inlet. Trimethylamine (CAS: 75-50-3, Sigma-Aldrich) and isopentylamine (CAS: 107-85-7, Sigma-Aldrich) were serially diluted to three concentrations in mineral oil. For sessions without odor, the air inlet was not switched. Sessions lasted 10 minutes, and the order of individual mice was randomized. One session per mouse was conducted each day, and the odor concentration was increased the following day. The apparatus was cleaned with 70% ethanol and vacuumed for at least 5 minutes after each session to clear the chambers of the introduced odor and any social cues left behind by the previous mouse.

Only behavior from the light chamber was recorded. Behavior videos were recorded and processed in ImageJ as previously described. Custom MATLAB code measuring the mouse’s location relative to its body midpoint was used to calculate the time spent in each chamber. The percent change was calculated as ((portion of time spent in the dark chamber after the UCMS protocol - portion of time spent in the dark chamber before the UCMS protocol) / portion of time spent in the dark chamber before the UCMS protocol) * 100. Statistical analysis was done using the rank-sum test and two-way analysis of variance (ANOVA). Values are given as mean ± SEM.

### Perfusion and Immunohistochemistry

Both control and stressed groups were anesthetized with a ketamine/xylazine solution (100 mg/10 kg) via intraperitoneal injection. The degree of anesthetization was verified via toe pinches before transcardial perfusion with 10 mL of phosphate-buffered saline, followed by 10 mL of 10% formalin. Brains from both control and stressed experimental groups were fixed overnight in 4 mL of formalin, then dehydrated in a 30% sucrose solution for at least 72 hours before sectioning. Brains were suspended in Optimal Cutting Temperature compound for cryosectioning with a Leica CM1800 cryostat at 40 µm thickness. Sections were stored at 4 °C in phosphate-buffered saline (PBS) until immunohistochemistry was performed.

For immunohistochemistry, 9-11 brain sections were collected from each animal. Sections were chosen based on the presence of the following regions of interest: glomerular and granule cell layer of the main olfactory bulb, the accessory olfactory bulb, the anterior olfactory nucleus, the anterior piriform cortex, and the medial amygdala. Sections from the piriform cortex were obtained from the anterior portion with the lateral olfactory tract still present. These sections were washed for 30 minutes in KPBS with 5-minute interval changes. The tissue was then blocked and permeabilized (KPBS with 0.3% Triton X-100 and 2% donkey serum) for 30 minutes. After washing once more in KPBS for 20 minutes with 5-minute interval changes, the tissue was incubated in rabbit anti-Iba1 (1:100, Fujifilm, catalog number 019-19,741) and placed on an orbital platform overnight at room temperature. The tissue was washed again in KPBS for 30 minutes with 5-minute intervals, then incubated in goat anti-GFAP (1:100, Abcam, catalog number AB_53554) on an orbital platform overnight at room temperature. After washing for 40 minutes in KPBS at 5-minute intervals, the tissue was incubated with secondary antibodies anti-RB-TRITC and anti-GT-FITC (1:500, Jackson ImmunoResearch, catalog number 705-095-003) in antibody diluent (1x KPBS, 0.02% sodium azide, 2% donkey serum). The tissue was washed for 30 minutes in KPBS with 5-minute change intervals and mounted on glass slides using ProLong Gold Antifade with DAPI (Invitrogen, catalog number P36931). Slides were stored at 4 °C until imaging.

### Image analysis for microglia and astrocytes

Images were obtained using a Zeiss LSM800 confocal microscope at 10x or 25x for particle or Sholl analysis. These images were captured semi-randomly as 10-20 plane Z-stacks (Z-interval = 1 µm), with the only requirements that they were taken within the previously listed regions of interest and that at least one full microglia and one full astrocyte were centered within the stack. For each region of interest (ROI), three images were taken from two different tissue sections for each animal. 3-6 animals were used for each experimental group.

For astrocytes, each 25x Z-stack image was processed in ImageJ. The image was split into separate color channels before being collapsed into a maximum-intensity projection. An image threshold was then set using the default parameters upon selecting the threshold type “Otsu.” After setting the threshold, Fiji’s “analyze particles” function was used with a pixel size of 15-infinity. The “Count” output was included in the analysis, and an unpaired t-test was used to compare astrocyte counts between groups when both groups were normally distributed (determined by the Lilliefors test) or a two-tailed rank-sum test when distributions were non-parametric. A total of 3-6 images were used per ROI for each experimental group. False discovery was corrected using the Benjamini-Hochberg procedure.

For microglia, 25x Z-stack images were processed in ImageJ. The image was collapsed into a maximum-intensity projection and then split into different color channels. From the maximum-intensity projection, lab members blinded to the experimental condition outlined random spatially isolated microglia using a freehand region of interest (ROI). The surrounding area was removed using ImageJ’s “clear outside” function, and an image threshold was set using the default parameters of the “Default” threshold type. An ROI was then indicated at the center of the soma, the single microglia mask was skeletonized using ImageJ’s “skeletonize” plugin, and a Sholl analysis was performed using the Fiji Sholl tool with the following parameters: start radius (from the center of the indicated ROI at the soma) = 3 µm, step size = 2 µm, and end radius = 50 µm. The number of identified intersections at each 2 µm step for each microglia, measured from each animal, was considered in the analysis. A one-way ANOVA was performed to compare the number of intersections at each 2 µm step across groups within each ROI, and the results were adjusted for false discovery using the Benjamini-Hochberg correction. For comparisons of the peak number of intersections, an unpaired t-test was used to compare measurements between groups when both groups were normally distributed (determined by the Lilliefors test) or a two-tailed rank-sum test when distributions were non-parametric. False discovery was corrected using the Benjamini-Hochberg procedure.

## Results

### Unpredictable stressors induce a depressive-like state in mice

We developed a standardized protocol (see *Methods*) consisting of six mild, but unpredictable stressors (UCMS) to induce a depressive-like state in adult male and female mice. Recent studies have also used a similar design to study habit formation in the context of stress (Giovanniello et al., 2025). Our panel of stressors and the unpredictability of their implementation resulted in a cumulative effect that gave rise to behavioral phenotypes that are characteristic of depression-like symptoms in rodent models (Mineur et al., 2007; Kreisel et al., 2014).

We began our study by first validating the efficacy of our approach. For each animal, we first collected baseline immobility measurements using two standard behavioral assays, the Forced Swim Test and the Tail Suspension Test (see *Methods*; **Figure 1A**). Animals were then randomly assigned to either a stress group that received the UCMS protocol or a control group that received equivalent daily handling. After four weeks, mice were then reassessed for immobility on both the Forced Swim Test and the Tail Suspension Test. During the initial Forced Swim Test, there was no difference in the immobility scores (defined as the fraction of time spent actively swimming) between the control and stressed groups (0.5700 ± 0.0648 in control animals, *n* = 11 and 0.6466 ± 0.0221 in stressed animals, *n* = 12; *p* = 0.4060, rank-sum test; **Figure 1B**; **Table 4**). However, following four weeks of UCMS, the stressed group showed an increase in their immobility score from baseline measurements (0.6466 ± 0.0221 before stressors and 0.8487 ± 0.0189 after stressors, *n* = 12; *p* = 4.8828x10^-4^, signed-rank test; Bonferroni corrected critical value = 0.0125; **Figure 1B**; **Table 4**), indicating less time attempting to swim in the chamber. A reduction in swimming time has been observed in other studies of learned helplessness and depression-like states in rodents (Yankelevitch-Yahav et al., 2015; Kraeuter et al., 2019). In the control group, there was no difference in the immobility scores following four weeks of daily handling with no stressors (0.5700 ± 0.0648 before stressors and 0.8487 ± 0.0189 after stressors, *n* = 11; *p* = 0.0322, signed-rank test; Bonferroni corrected critical value = 0.0125; **Figure 1B**; **Table 4**).

**Table 4.** Figure 1 statistics.

| Behavioral Test | Comparison | p-value | z-statistic | n | Test |
| --- | --- | --- | --- | --- | --- |
| Forced Swim Test | Control-Control before and after stressors | 0.0329 | 2.1339 | 11 control<br>11 stressed | signed-rank |
|  | **Stressed-Stressed before and after stressors** | **$4.8828 \times 10^{-4}$** | **3.0594** | **12 control**<br>**12 stressed** | **signed-rank** |
|  | Control-Stressed before stressors | 0.4060 | 0.8309 | 11 control<br>12 stressed | rank-sum |
|  | Control-Stressed after stressors | 0.0151 | 2.4311 | 11 control<br>12 stressed | rank-sum |
| Tail Suspension Test | Control-Control before and after stressors | 0.2860 | 1.0669 | 11 control<br>11 stressed | signed-rank |
|  | **Stressed-Stressed before and after stressors** | **$4.8828 \times 10^{-4}$** | **3.0594** | **12 control**<br>**12 stressed** | **signed-rank** |
|  | Control-Stressed before stressors | 0.8294 | -0.2154 | 11 control<br>12 stressed | rank-sum |
|  | Control-Stressed after stressors | 0.0337 | 2.1233 | 11 control<br>12 stressed | rank-sum |
| Open Field Test | Control-Control before and after stressors | 0.7002 | -0.4446 | 11 control<br>11 stressed | signed-rank |
|  | Stressed-Stressed before and after stressors | 0.0923 | -1.7258 | 12 control<br>12 stressed | signed-rank |
|  | Control-Stressed before stressors | 0.4060 | -0.8309 | 11 control<br>12 stressed | rank-sum |
|  | Control-Stressed after stressors | 0.2301 | -1.2001 | 11 control<br>12 stressed | rank-sum |

The Tail Suspension Test similarly showed no difference between groups in the amount of time spent immobile prior to UCMS (0.4928 ± 0.0305 in control animals, *n* = 11, and 0.4932 ± 0.0409 in stressed animals, *n* = 12; *p* = 0.8294, rank-sum test; **Figure 1C**; **Table 4**). Following UCMS, the stressed group again showed reduced time struggling (0.4932 ± 0.0409 before stressors and 0.7908 ± 0.0428 after stressors, *n* = 12; *p* = 4.8828 x10^-4^, signed-rank test; Bonferroni corrected critical value = 0.0125; **Figure 1C**; **Table 4**), while there was no difference in the control group after daily handling (0. 4928 ± 0.0305 before stressors and 0.5618 ± 0.0797 after stressors, *n* = 11; *p* = 0.3203, signed-rank test; Bonferroni corrected critical value = 0.0125; **Figure 1C**; **Table 4**).

As a final test to ensure that UCMS did not result in generalized motor deficits, which could manifest as reduced struggle behavior, we measured locomotor activity in an open-field arena for each mouse. In both groups, after the UCMS protocol, there was no difference in their normalized mean velocity and total movement as they traversed the arena (10.8299 ± 0.9064 cm/s before stressors and 11.5830 ± 1.2488 cm/s after stressors in the control group, *p* = 0.7002, *n* = 11, signed-rank test; 11.4210 ± 0.6032 cm/s before stressors and 13.6825 ± 0.7658 cm/s after stressors in the stressed group, *p* = 0.0923, *n* = 12, signed-rank test; **Figure 1D**; **Table 4**). Because the observed behavioral alterations occurred in the absence of motor impairments, this suggests that UCMS selectively alters affective state rather than general activity levels (Mineur et al., 2006; Antoniuk et al., 2019). These results demonstrate that our four-week paradigm of daily unpredictable mild stressors is sufficient to induce depressive-like behaviors in mice, resulting in a reduction of self-preservation behaviors.

### Avoidance of aversive olfactory stimuli is increased following exposure to stressors

We next wanted to understand how depressive-like states may influence both tolerance and aversion of olfactory stimuli. We hypothesized that the depressive-like state conferred by UCMS would render mice less tolerant of aversive olfactory stimuli.

We devised a behavioral paradigm comprising two chambers (see *Methods*) connected by a tunnel. One chamber was transparent and illuminated, while the other was made completely opaque and darkened. Mice were allowed to freely explore the apparatus for five minutes, after which the darkened chamber was odorized using air inlets. Odorants were delivered to a carrier stream so that air was constantly flowing to the dark chamber for the duration of the experiment (**Figure 2A**). Earlier work characterizing mouse behavior preferences for illuminated and darkened chambers revealed that mice spend most of their time in darkened chambers (Costall et al., 1989). This allowed us to ask how odor quality and aversion can be balanced with the relative “safety” of the dark environment.

**Figure 2.**
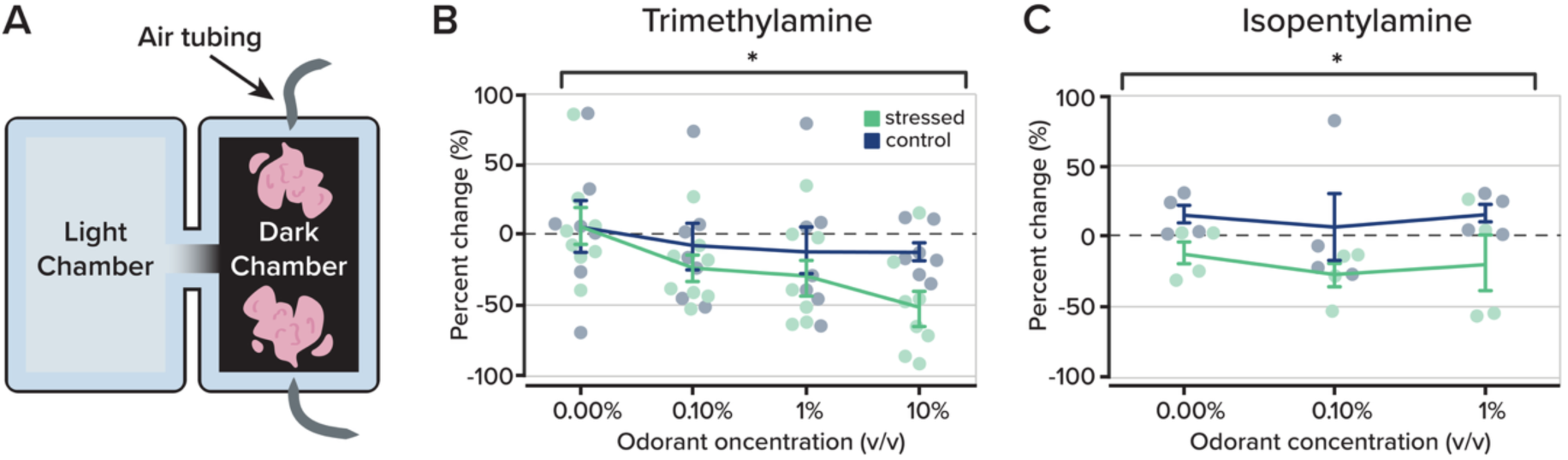
Changes in olfactory-guided behavior in an odorized Light/Dark Box Test. **A.** Odorized Light/Dark Box apparatus set up. Odorants trimethylamine and isopentylamine were introduced in the dark chamber at different concentrations using v/v dilutions (0%, 0.1%, 1%, 10%). **B.** Behavioral response to introduction of odorant trimethylamine plotted as percent change (%) of the time spent in the dark chamber from before and after exposure to the UCMS protocol between the control (*n* = 7) and stressed (*n* = 8) groups. A difference was found across all odorant concentrations using a two-way ANOVA test with repeated measures: F_(3,1)_ = 5.7300, *p* = 0.0020. A rank-sum test revealed a difference between groups at 10% concentration, *p* = 0.0289. **C.** As in *part B*, behavioral responses to the odorant isopentylamine are plotted to visualize comparisons between the control (*n* = 4) and stressed (*n* = 4) groups. A two-way ANOVA with repeated measures revealed a difference between experimental groups across all concentrations: F_(2,1)_ = 8.8300, *p* = 0.0250. Data are displayed as mean ± SEM. Individual dots represent experimental replicates.

We tested two separate odorants known to have aversive qualities to mice, trimethylamine and isopentylamine (Belonio et al., 2025; Root et al., 2014), but that are not associated with predator cues. Mice were tested across a range of concentrations for both odorants (0% - 10% v/v for trimethylamine and 0% - 1% v/v for isopentylamine). The time spent in the darkened chamber was then compared with the pre-odor period and expressed as a percent change. For both odorants, we found that the stressed group spent less time in the odorized chamber than the control group (**Figure 2B,C**). At the highest concentration (10% v/v) of trimethylamine, the stressed group spent 51.3260 ± 15.9051% (*n* = 8) less time in the darkened odorized chamber compared to 12.6859 ± 6.7897% (*n* = 7) less time in the control group (*p* = 0.0289; *z*-statistic = 2.1410; rank-sum test). When we considered behavioral aversion across all odorant concentrations, the stressed group showed a stronger avoidance of the odorized compartment than the control group (*p* = 0.0020; F_(3,1)_ = 5.7300; two-way ANOVA; **Figure 2B**). A similar relationship was observed with the odorant isopentylamine: the stressed group spent less time in the odorized compartment (*p* = 0.0250; F_(2,1)_ = 8.8300; two-way ANOVA; **Figure 2B**).

Together, our results indicate that, in our stress induced model of a depression-like state, mice show increased sensitivity and decreased tolerance to aversive odorants. This was apparent as stressed mice were more willing to leave the relative safety of the darkened chamber when it became odorized.

### Neuroinflammatory markers associated with chronic stress

Neuroinflammation is a biomarker of chronic stress and is also associated with depression (Troubat et al., 2021; Guo et al., 2023). Among the primary cellular mediators of neuroinflammation are astrocytes, which become reactive under pathological conditions and upregulate glial fibrillary acidic protein (GFAP), a hallmark of astrogliosis. To assess whether UCMS elicited neuroinflammatory responses in olfactory and limbic brain regions, we examined astrocyte activation across multiple areas known to process odor information and emotional valence.

Following behavioral testing, brain tissue was stained for GFAP (see *Methods*; **Figure 3A**). We quantified GFAP-positive astrocytes as a readout of glial activation in both primary sensory and higher-order olfactory regions, as well as in the medial amygdala, a region implicated in both olfactory processing and emotional regulation (Swanson & Petrovich, 1998; Janak & Tye, 2015). Our initial analysis revealed that across all brain areas, there was a difference in the number of GFAP-positive astrocytes (F(5,25) = 50.3659, *p* = 0.3737 x 10^-19^, two-way ANOVA); however, there was no difference between experimental groups (F(1,25) = 0.3642, *p* = 0.5490, two-way ANOVA).

**Figure 3.**
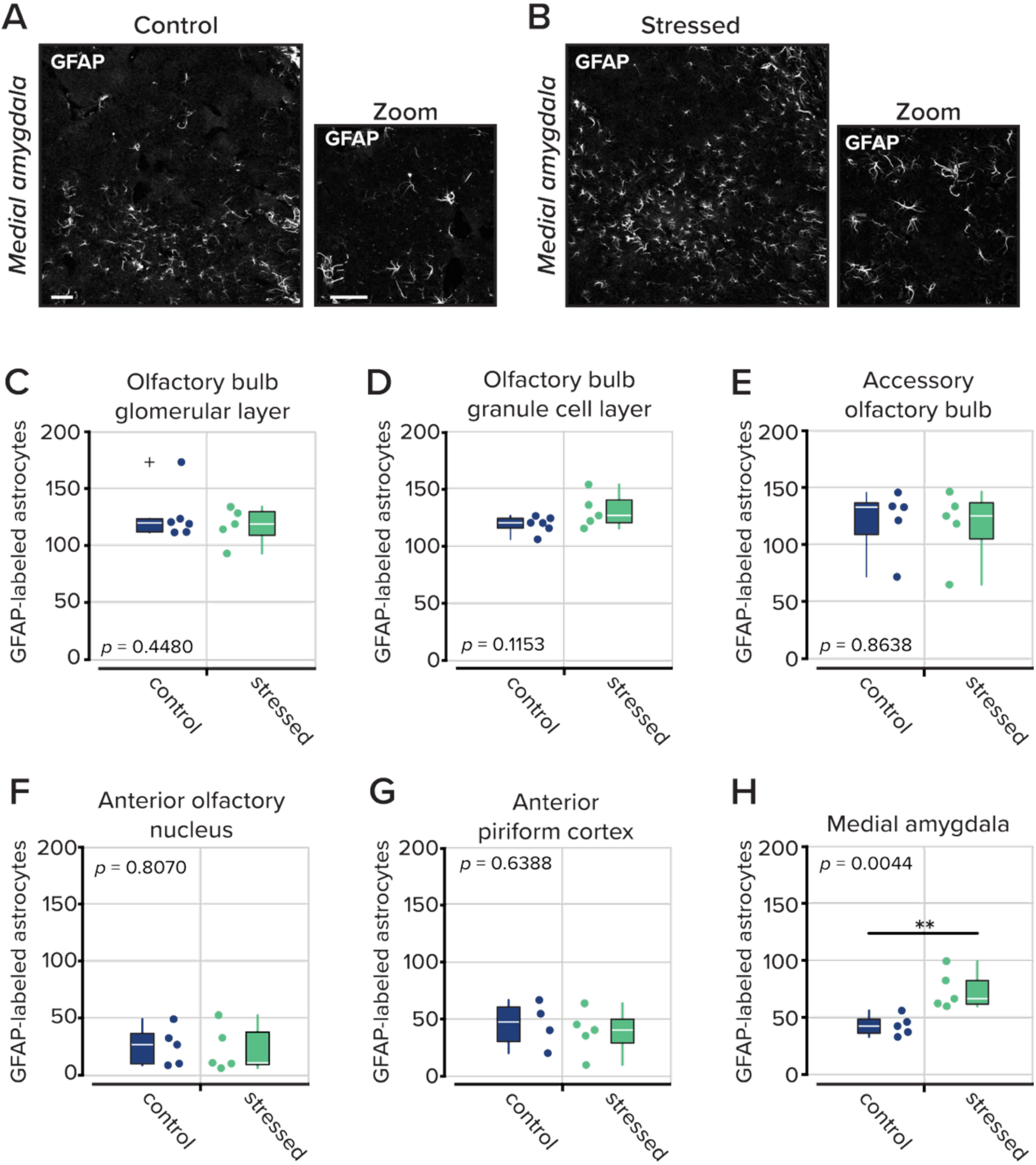
GFAP-labeled astrocytes throughout the olfactory system. **A.** Representative images of astrocytes labeled with GFAP in the medial amygdala at 10x (*left*) and 40x (*right*). Scale bar denotes 50 µm. **B.** Same as *part A* for an animal that underwent the UCMS protocol. **C-H.** The total number of GFAP-labeled astrocytes counted in different regions of interest: the olfactory bulb glomerular layer, olfactory bulb granule cell layer, accessory olfactory bulb, anterior olfactory nucleus, anterior piriform cortex, and medial amygdala. In the medial amygdala, a difference between groups was revealed (*p* = 0.0044). Dots represent experimental replicates. The white line in the box represents the median, the box edges indicate the 25th and 75th percentiles, respectively. The whiskers extend to the most extreme data points. Outliers are plotted individually using the ‘+’ marker symbol. *n* = 4-6 animals per group, three to six images were analyzed per region of interest in each animal.

We then considered each brain area individually. At the earliest stages of sensory processing, in the olfactory and accessory olfactory bulbs, we found no difference in the number of activated astrocytes (**Figure 3C-H**) between the control and stressed cohorts (**Table 5**). Similarly, in the olfactory cortex, the anterior piriform cortex, and the anterior olfactory nucleus, there was also no difference in the total number of GFAP-positive astrocytes. However, in the medial amygdala, which receives olfactory input and encodes affective state, we found more activated astrocytes in the stressed group than in their non-stressed counterparts (45.2333 ± 3.0816 astrocytes per imaging field in the control group and 76.8333 ± 7.4375 in the stressed group. (*n* = 5 mice (three to six imaging fields from each animal); *p* = 0.0044; rank-sum test). Our findings indicate that repeated stressors drive neuroinflammation, as evidenced by activated astrocytes at loci receiving convergent input from both affective and sensory areas.

**Table 5.** Figure 3 statistics.

| Region | Control mean ± SEM | Stressed mean ± SEM | Statistic | $n$ | $p$-value | Test | Sig. |
| --- | --- | --- | --- | --- | --- | --- | --- |
| Olfactory bulb glomerular layer | 123.0278 ± 9.2601 | 114.3667 ± 6.8899 | $z = 0.183$ | 6 control<br>5 stressed | 0.9307 | rank-sum | ns |
| Olfactory bulb granule cell layer | 112.9444 ± 2.9252 | 124.6333 ± 6.5091 | $t = -1.743$ | 6 control<br>5 stressed | 0.1153 | t-test | ns |
| Accessory olfactory bulb | 116.9833 ± 12.5745 | 109.5333 ± 13.0697 | $t = 0.411$ | 5 control<br>5 stressed | 0.6920 | t-test | ns |
| Anterior olfactory nucleus | 26.2000 ± 7.4697 | 23.2833 ± 8.8129 | $t = 0.252$ | 5 control<br>5 stressed | 0.8070 | t-test | ns |
| Anterior piriform cortex | 43.5000 ± 9.9671 | 37.0333 ± 8.6957 | $t = 0.113$ | 4 control<br>5 stressed | 0.9127 | t-test | ns |
| **Medial amygdala** | **45.2333 ± 3.0816** | **76.8333 ± 7.4375** | **$t = -3.925$** | **5 control**<br>**5 stressed** | **0.0044** | **t-test** | **\*\* (FDR sig)** |

Values are mean ± SEM. Normality was tested with the Lilliefors test; an unpaired t-test was applied where both groups passed normality, and a rank-sum test otherwise. $p$ -values FDR-corrected (Benjamini-Hochberg) across 6 areas. \*\* $p < 0.01$ (FDR sig denotes significance after correction).

### Stress induces microglial remodeling in the olfactory system

In many neurodegenerative disease models, microglia decrease the complexity and size of their processes (Davalos et al., 2005; Qin et al., 2023; Fyke et al., 2024). However, under chronic inflammatory conditions, microglia can develop hypertrophic morphologies (Hains and Waxman, 2006; Loane et al., 2014; Calcia et al., 2016). To test whether our model of a stress-induced depression-like state was sufficient to induce microglial activation and morphological changes, we immunostained the same animals for ionized calcium-binding adaptor molecule 1 (Iba1), a protein enriched in microglia and other macrophages (Ito et al., 1998). From images of the same olfactory areas previously described, we isolated individual microglia, measured process complexity using Sholl analysis (see *Methods*), and counted path intersections at each concentric ring centered on the microglial soma.

From four to six mice in each group, we selected two images from each area. In each image, two microglia were selected by an experimenter who was blind to the animal condition for further analysis. Our results revealed microglial remodeling in two key olfactory structures: the anterior olfactory nucleus and anterior piriform cortex. In both regions, microglia from UCMS-treated mice exhibited more complex and extended processes than controls (in the anterior olfactory nucleus: *p* = 0.0130; in the anterior piriform cortex: *p* = 0.0001; one-way ANOVA; **Figure 4F,G**; **Table 6**), indicative of hypertrophic remodeling. Interestingly, no differences were observed in the olfactory bulb, accessory olfactory bulb, or medial amygdala, suggesting region-specific vulnerability or temporal dynamics of microglial activation in response to chronic stress.

**Figure 4.**
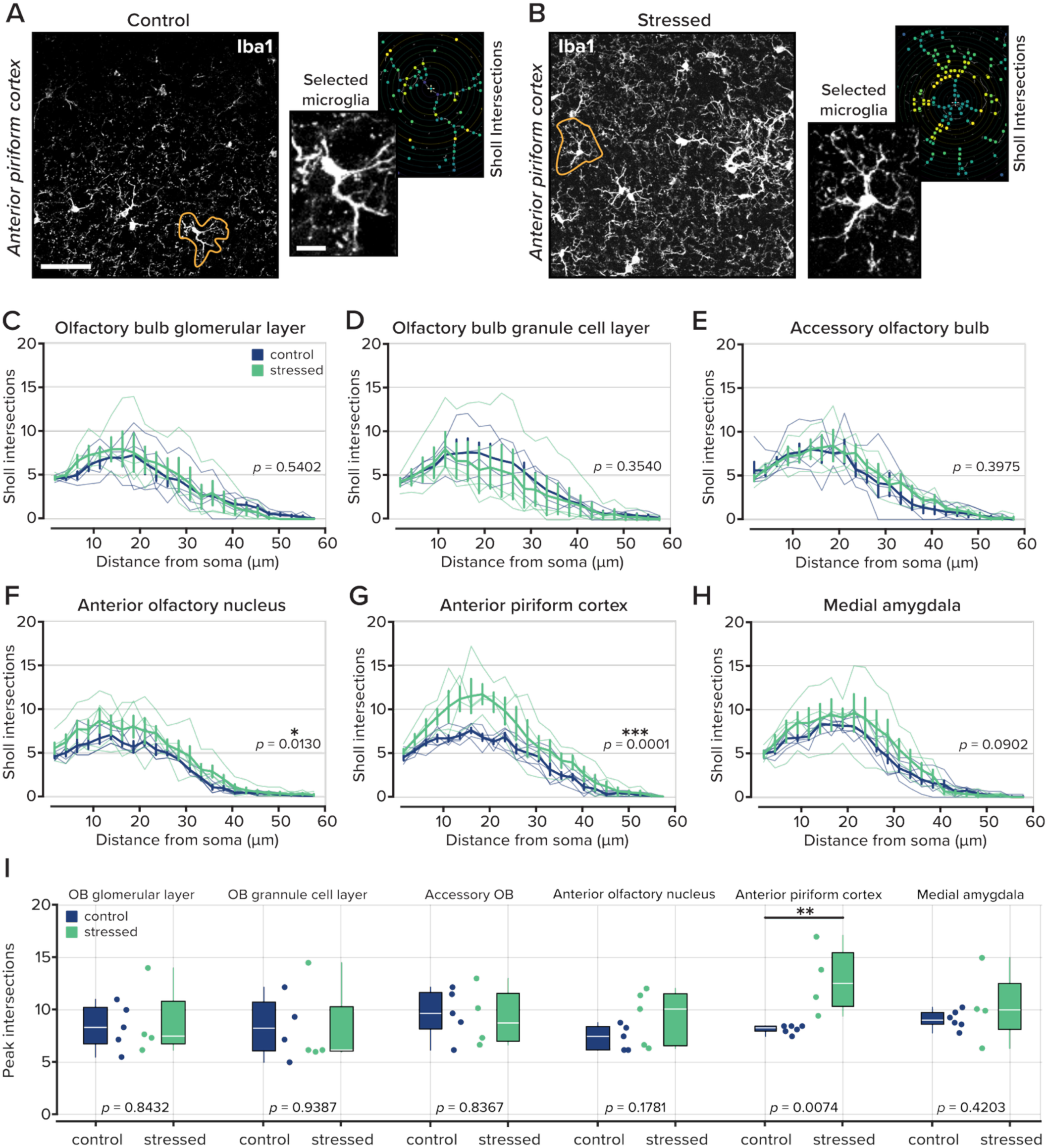
Iba1-labeled microglia in olfactory brain areas. **A.** Representative images of microglia labeled with Iba1 in the glomerular layer of the olfactory bulb at 25x. The encircled cell is the cell used for analysis. Scale bar denotes 50 µm. Right, an expanded version of the selected microglia and Sholl intersections. Inset scalebar is 10 µm. **B**. Same as *part A* for an animal that underwent the UCMS protocol. **C-H.** The number of microglial processes, represented as Sholl intersections, by distance away from the cell soma, in different regions of interest: the olfactory bulb glomerular layer, olfactory bulb granule cell layer, accessory olfactory bulb, anterior olfactory nucleus, anterior piriform cortex, and medial amygdala. Statistical tests were performed with a one-way ANOVA. Data are displayed as mean ± SEM, with individual animal averages shown as thin colored lines. *n* = 4-6 animals per group, 4-8 microglia for each animal. **I**. The mean peak value of Sholl intersections for each animal in all six brain regions.

**Table 6.**
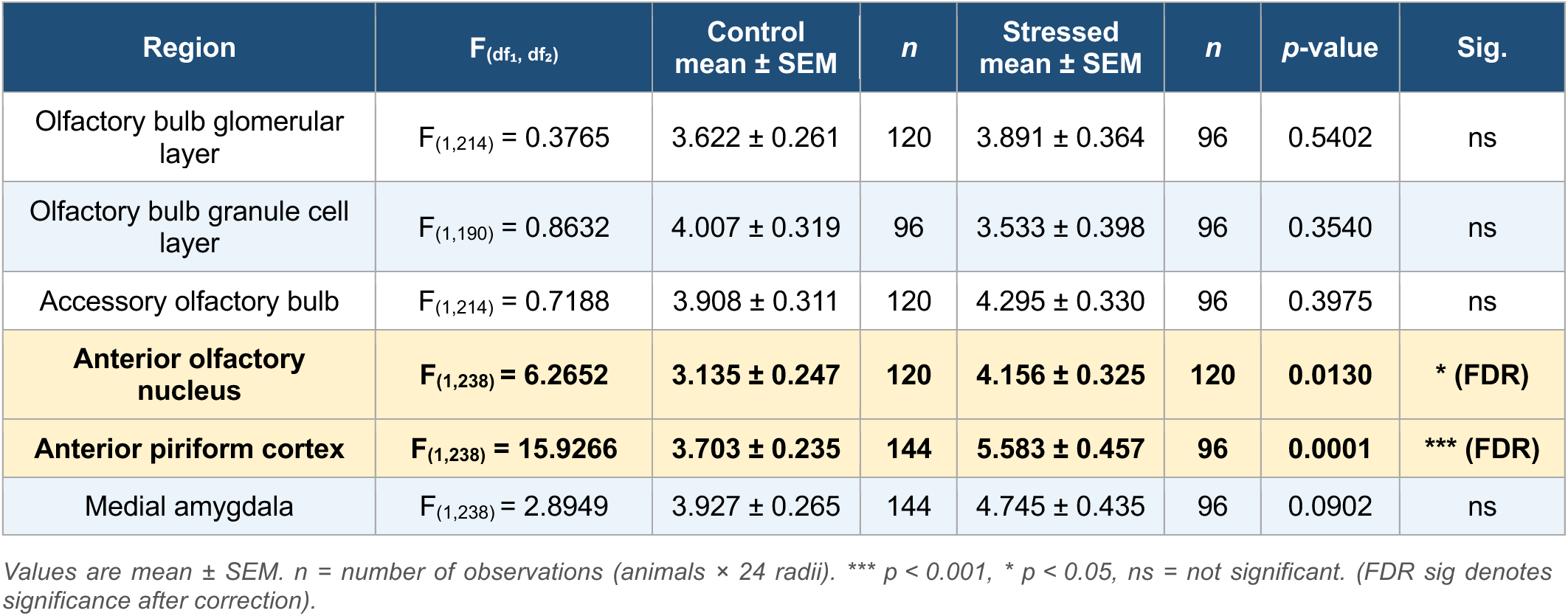
Figure 4C-H statistics.

| Region | $F_{(df_1, df_2)}$ | Control mean ± SEM | $n$ | Stressed mean ± SEM | $n$ | $p$-value | Sig. |
| --- | --- | --- | --- | --- | --- | --- | --- |
| Olfactory bulb glomerular layer | $F_{(1,214)} = 0.3765$ | $3.622 \pm 0.261$ | 120 | $3.891 \pm 0.364$ | 96 | 0.5402 | ns |
| Olfactory bulb granule cell layer | $F_{(1,190)} = 0.8632$ | $4.007 \pm 0.319$ | 96 | $3.533 \pm 0.398$ | 96 | 0.3540 | ns |
| Accessory olfactory bulb | $F_{(1,214)} = 0.7188$ | $3.908 \pm 0.311$ | 120 | $4.295 \pm 0.330$ | 96 | 0.3975 | ns |
| **Anterior olfactory nucleus** | **$F_{(1,238)} = 6.2652$** | **$3.135 \pm 0.247$** | **120** | **$4.156 \pm 0.325$** | **120** | **0.0130** | **\* (FDR)** |
| **Anterior piriform cortex** | **$F_{(1,238)} = 15.9266$** | **$3.703 \pm 0.235$** | **144** | **$5.583 \pm 0.457$** | **96** | **0.0001** | **\*\*\* (FDR)** |
| Medial amygdala | $F_{(1,238)} = 2.8949$ | $3.927 \pm 0.265$ | 144 | $4.745 \pm 0.435$ | 96 | 0.0902 | ns |

Values are mean ± SEM. $n$ = number of observations (animals × 24 radii). \*\*\* $p < 0.001$ , \* $p < 0.05$ , ns = not significant. (FDR sig denotes significance after correction).

We also considered the peak value of Sholl intersections across areas and groups. A two-way ANOVA revealed no difference between groups across all brain areas (F_(1,24)_ = 3.1767; *p* = 0.0689). However, when we considered each brain area individually, there was an increase in the peak value of the microglia Sholl intersections in the anterior piriform cortex (8.0278 ± 0.1634 intersections in control animals, *n* = 6 animals and 12.8750 ± 1.7029 intersections in stressed animals, *n* = 4 animals; *p* = 0.0074; unpaired t-test, FDR corrected significance; **Figure 6I**; **Table 7**). No other brain areas showed differences in Sholl intersection peak values.

**Table 7.** Figure 4H statistics.

| Region | Control mean ± SEM | Stressed mean ± SEM | Statistic | $n$ | $p$-value | Test | Sig. |
| --- | --- | --- | --- | --- | --- | --- | --- |
| Olfactory bulb glomerular layer | $8.400 \pm 0.9812$ | $8.7917 \pm 1.7656$ | $t = -0.2052$ | 5 control<br>4 stressed | 0.8432 | t-test | ns |
| Olfactory bulb granule cell layer | $8.4167 \pm 1.5313$ | $8.2083 \pm 2.0976$ | $z = 0.2887$ | 4 control<br>4 stressed | 0.8000 | rank-sum | ns |
| Accessory olfactory bulb | $9.6667 \pm 1.0620$ | $9.2917 \pm 1.4504$ | $t = 0.2139$ | 5 control<br>4 stressed | 0.8367 | t-test | ns |
| Anterior olfactory nucleus | $7.2333 \pm 0.5467$ | $9.2000 \pm 1.2150$ | $t = -1.4762$ | 5 control<br>5 stressed | 0.1781 | t-test | ns |
| **Anterior piriform cortex** | **$8.0278 \pm 0.1634$** | **$12.8750 \pm 1.7029$** | **$t = -3.5598$** | **6 control**<br>**4 stressed** | **0.0074** | **t-test** | **\*\* (FDR sig)** |
| Medial amygdala | $8.9722 \pm 0.3637$ | $10.2500 \pm 1.8137$ | $t = -0.8495$ | 6 control<br>4 stressed | 0.4203 | t-test | ns |

Values are mean ± SEM. Normality was tested with the Lilliefors test; an unpaired t-test was applied where both groups passed normality, and a rank-sum test otherwise. $p$ -values FDR-corrected (Benjamini-Hochberg) across 6 areas. \*\* $p < 0.01$ , ns = not significant. (FDR sig denotes significance after correction).

These findings contrast with our earlier observations of astrocyte reactivity, which was largely restricted to the medial amygdala. While astrocytic changes appeared localized, microglial remodeling was more pronounced in cortical olfactory processing regions, implicating microglia as sensitive responders to stress-related neuroinflammation. Together, these data suggest that UCMS elicits differential patterns of glial activation, with microglia in the anterior olfactory nucleus and anterior piriform cortex exhibiting robust morphological changes consistent with heightened immune surveillance and potential modulation of local circuit function.

### Microglial complexity and astrocyte density are independently regulated following chronic stress

To examine whether microglial morphological complexity and astrocyte density were co-regulated within individual animals, we performed Pearson correlations between peak Sholl intersections and astrocyte counts across matched animals in each brain area. When pooled across groups, correlations were uniformly weak (|ρ| ≤ 0.34; **Figure 5A-F**) and non-significant in all six areas examined (all *p* > 0.36, FDR-corrected; **Table 8**). These results indicate that microglial morphological complexity and astrocyte density vary independently across animals, suggesting that stress-induced changes in these two cell populations are driven by distinct rather than shared regulatory mechanisms.

**Figure 5.**
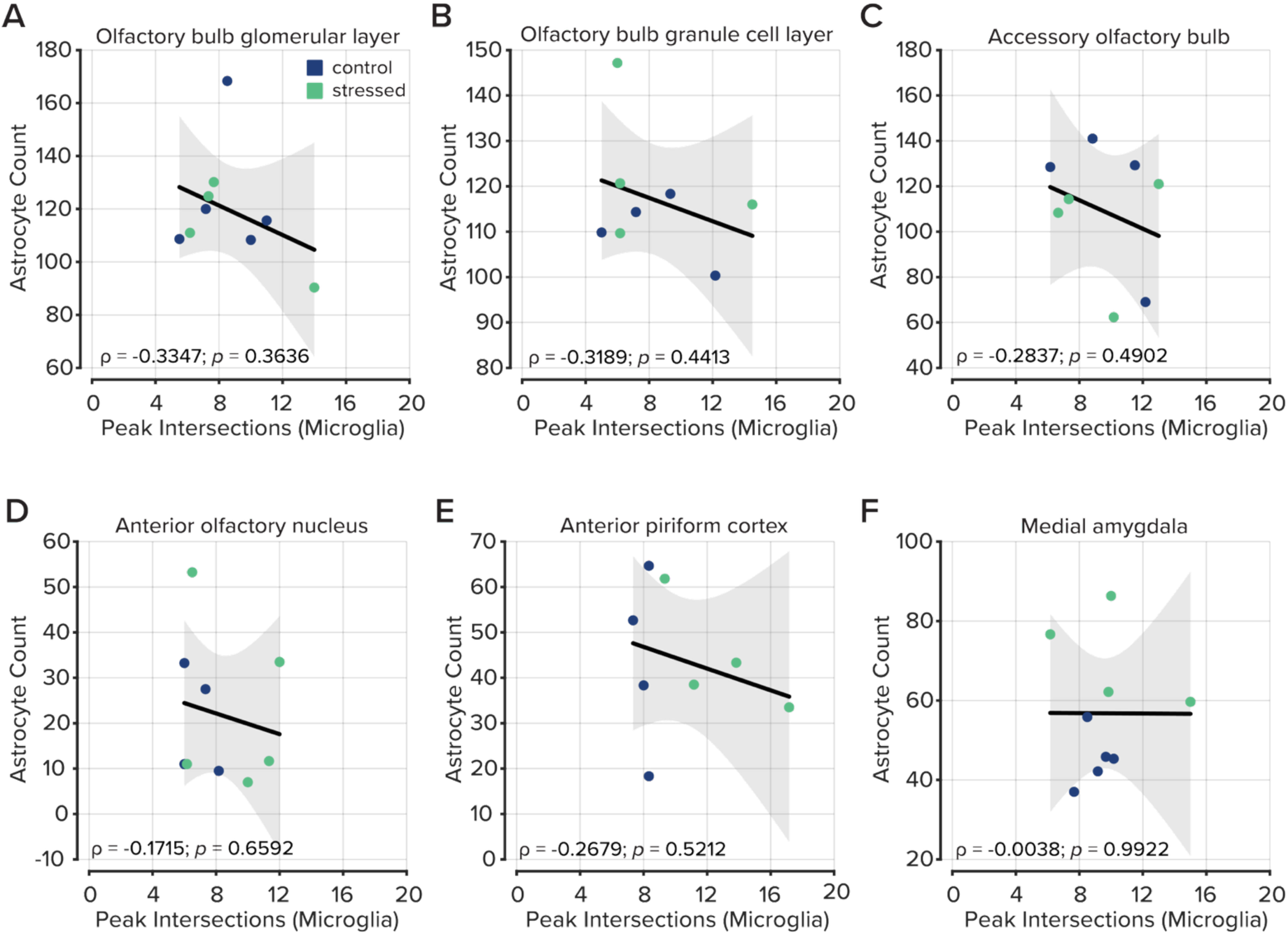
Correlation analysis between astrocyte counts and microglial morphology. **A-F.** The total number of astrocytes plotted by the peak value of Sholl intersections per animal in each brain region. No regions show a relationship between astrocyte counts and microglial morphology.

**Table 8.**
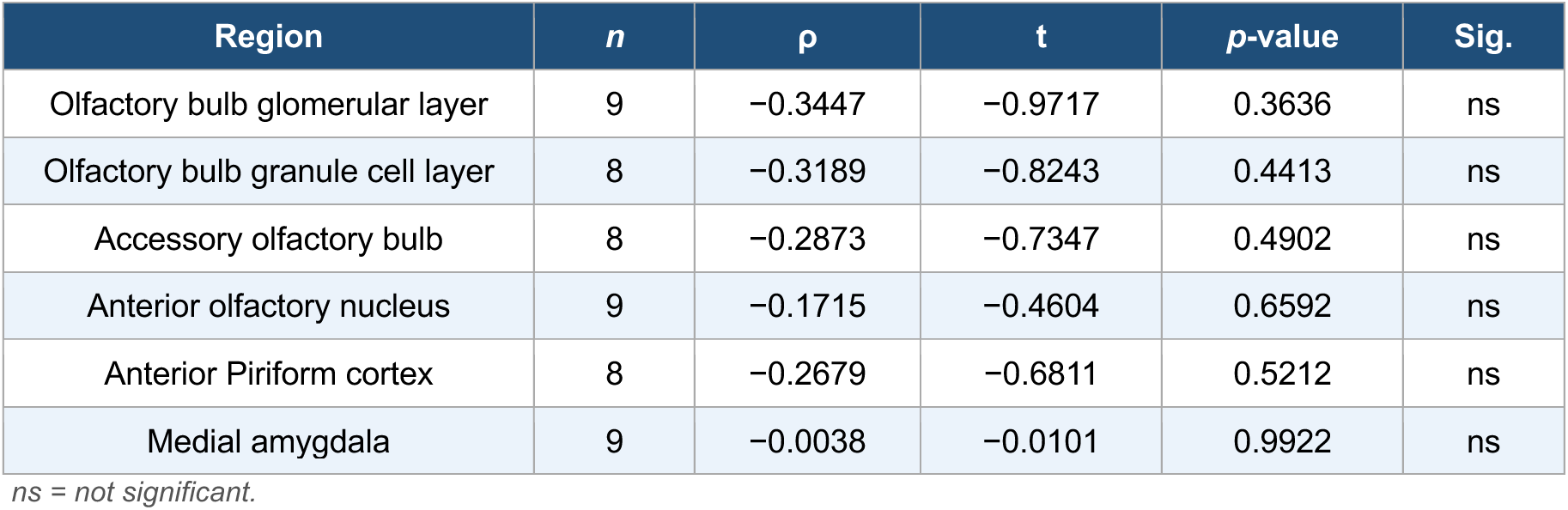
Figure 5 statistics.

### Behavioral olfactory avoidance is related to neuroinflammation in key olfactory areas

In the final part of our study, we asked whether behavioral olfactory avoidance was associated with the degree of neuroinflammation in each of the olfactory areas we previously analyzed. We identified three areas that demonstrated a relationship between behavioral olfactory avoidance and glial activation: the medial amygdala, the anterior olfactory nucleus, and the anterior piriform cortex.

In the medial amygdala, we compared changes in time spent in the dark chamber after odorization with 10% trimethylamine with the total number of astrocytes. Here, we selected the medial amygdala because we previously observed increased astrocyte counts (**Figure 3H**). In this analysis, we combined the two cohorts of mice due to individual variability in behavior and astrogliosis. Our analysis revealed a relationship between these variables (ρ = -0.7050, t = -2.8118, *p* = 0.0228, *n* = 10; **Figure 6A**; **Figure 6-1**); however, after false discovery rate correction using the Benjamini-Hochberg procedure, the relationship was not considered significant. In the other areas we included in our analysis, we found no relationship between the number of astroglia and behavioral outcomes (**Figure 6-1**; **Figure 6-2**), thereby indicating that astrogliosis in olfactory processing areas does not play a role in olfactory avoidance.

**Figure 6.**
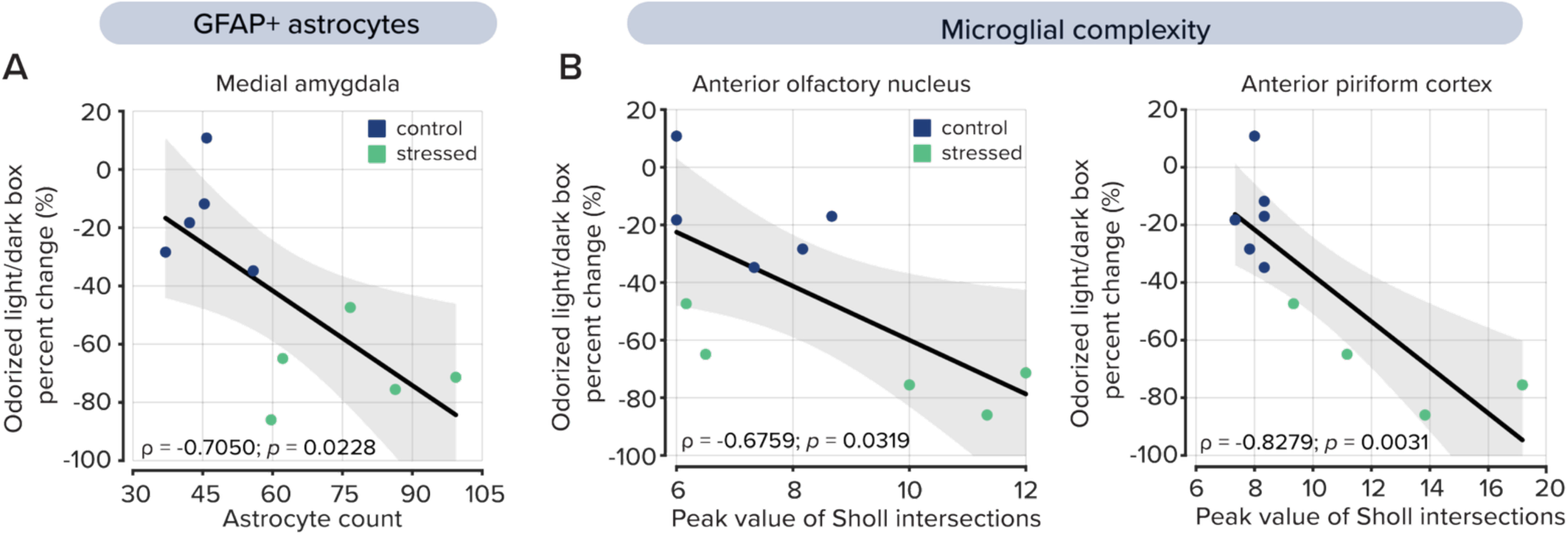
Correlation analysis between inflammatory response and behavior in the Odorized light/dark box test. **A.** The total number of astrocytes in the medial amygdala plotted by the percent change of time spent in the dark chamber with the odorant trimethylamine. Plotted values represent each animal’s results from the behavioral assessment and IHC imaging. *p*-values were determined by calculating the t-value from the linear regression, *p* = 0.0228. Summary statistics across all brain regions are shown in the extended data Figure 6-2, and correlation plots are shown in the extended data Figure 6-1. **B.** The peak number of intersections identified by Sholl analysis in the microglia, as measured by the percent change in time spent in the dark chamber with the odorant trimethylamine in the anterior olfactory nucleus (*left*, *p* = 0.0319) and anterior piriform cortex (*right*, *p* = 0.0031). Summary statistics across all brain regions are shown in the extended data Figure 6-3, and correlation plots are shown in the extended data Figure 6-1.

We also considered microglial complexity as a driver of olfactory avoidance. In this analysis, we focused on the anterior olfactory nucleus and the anterior piriform cortex because they showed increased microglial complexity following UCMS (**Figure 4F,G**). Here, we compared the maximum number of Sholl intersections for all microglia in the anterior olfactory nucleus and the anterior piriform cortex of each mouse with behavioral avoidance of 10% trimethylamine.

In the anterior olfactory nucleus, we found a relationship between microglial morphological complexity and olfactory avoidance (ρ = -0.6759, t = −2.5942, *p* = 0.0319; *n* = 10; **Figure 6B**, *left*; **Figure 6-1**); however, the relationship did not withstand false discovery rate correction. We also found a relationship between behavioral avoidance and microglial complexity in the anterior piriform cortex (ρ = -0.8279, t = - 4.1751, *p* = 0.0031; *n* = 10; **Figure 5B**; **Figure 6-1**), which remained significant after false discovery rate correction. In all other regions, we found no relationship between microglial complexity and olfactory avoidance (**Figure 6-1**; **Figure 6-3**).

Together, these findings suggest that olfactory avoidance behavior is selectively associated with glial activation in distinct brain regions involved in olfactory processing and emotional salience. The absence of similar correlations in other olfactory areas highlights the regional specificity of these effects and suggests that glial activity in the anterior piriform cortex may uniquely contribute to the neural mechanisms underlying stress-induced changes in olfactory perception and avoidance.

## Discussion

In humans, chronic stress and depression are known to alter olfactory perception, tolerance, and aversion (Lombion-Pouthier et al., 2006; Atanasova et al., 2010; Athanassi et al., 2021). Clinical studies have shown that individuals with major depressive disorder exhibit reduced olfactory sensitivity, decreased hedonic ratings of pleasant odors, and heightened aversion to unpleasant ones (Pause et al., 2001; Lombion-Pouthier et al., 2006; Croy et al., 2014). These alterations are not merely perceptual deficits but are thought to reflect broader disruptions in how affective salience is assigned to sensory stimuli. For example, depressed patients often show diminished approach behavior to rewarding odors and increased withdrawal from aversive odors (Atanasova et al., 2010; Rochet et al., 2018).

Our findings recapitulate depression-like-induced olfactory sensitivity in a rodent model. UCMS-treated mice displayed a depressive-like state, characterized by increased immobility in the Forced Swim Test and Tail Suspension Test, behaviors commonly used to model learned helplessness and depression-like states (Porsolt et al., 1977; Steru et al., 1985; Willner, 2005). UCMS-treated mice also showed an increased avoidance of odorized environments, even when those environments were otherwise preferred (i.e., dark and enclosed; Costall et al., 1989). This suggests that depressive-like states in mice, like in humans, sensitize individuals to olfactory threats and reduce tolerance for aversive stimuli. Given the strong anatomical and functional links between olfactory and limbic circuits (Soudry et al., 2011; Yao et al., 2020), these data further support the hypothesis that altered olfactory processing in depression-like states arises from disrupted integration between sensory input and emotional valence.

Our findings extend beyond behavioral phenotyping by identifying distinct patterns of neuroinflammatory activation in olfactory and limbic brain regions following UCMS. Astrocytic activation, as measured by GFAP expression, was selectively elevated in the medial amygdala, a region implicated in the affective appraisal of sensory stimuli and the integration of olfactory and emotional information (Swanson & Petrovich, 1998; Janak & Tye, 2015). In contrast, microglial remodeling was observed in cortical areas including the anterior olfactory nucleus and anterior piriform cortex, where increased process complexity is a known hallmark of chronic stress-induced microglial activation (Kreisel et al., 2014; Calcia et al., 2016). These cell-type- and region-specific differences suggest that astrocytes and microglia may respond to chronic stress via distinct mechanisms that differentially affect sensory and emotional networks (Liddelow et al., 2017; Yirmiya et al., 2015).

In the final part of our study, we asked whether glial activation in these regions was associated with olfactory avoidance behavior. The number of GFAP-positive astrocytes in the medial amygdala correlated with increased avoidance of aversive odorants; however, when considering all six brain areas, these results could be attributed to false discovery. While astrogliosis in the medial amygdala may alter affective salience encoding and influence behavioral responses to environmental threats (Tynan et al., 2013; Seo et al., 2016), more studies are necessary to clarify its role. Conversely, microglial complexity in the anterior piriform cortex was correlated with odor avoidance, possibly implicating immune-mediated modulation of cortical sensory circuits in the processing of aversive olfactory stimuli (Chen et al., 2016; Füzesi et al., 2016). The absence of similar relationships in other olfactory areas supports the idea of regional specificity in how neuroinflammation affects sensory-driven behavior.

In the present study, immunohistochemical sections were sampled from the anterior piriform cortex; however, the piriform cortex is not structurally or functionally homogeneous. The anterior and posterior subdivisions differ substantially in their connectivity, with the anterior piriform cortex receiving dense direct input from the olfactory bulb and projecting primarily to other olfactory and prefrontal regions, while the posterior piriform cortex receives comparatively less direct olfactory input and instead maintains stronger reciprocal connections with associative and limbic areas including the amygdala and entorhinal cortex (Haberly, 2001; Gottfried, 2010). These differences in input-output organization suggest that the two subdivisions may serve distinct functional roles in odor processing, with the anterior piriform cortex more closely associated with early sensory representation and the posterior piriform cortex more involved in the integration of olfactory information with contextual and affective signals (Howard et al., 2009; Zelano et al., 2011). Given that chronic stress differentially affected glial populations across olfactory brain regions in the present study, it is plausible that microglial and astrocytic changes within the piriform cortex itself are not uniform along its anterior-posterior axis. Future studies examining glial morphology across both anterior and posterior subdivisions of the piriform cortex would help determine whether the effects we observe are specific to early sensory processing nodes or extend into integrative regions more closely connected with limbic and affective systems.

Together, our findings support a model in which chronic stress alters odor perception and behavioral reactivity by remodeling glia in cortical sensory areas. While previous work has largely focused on neuroinflammation in classic limbic structures such as the hippocampus and prefrontal cortex (Duman et al., 2016; Wohleb et al., 2018), our data highlight the olfactory system as a novel site of stress-sensitive neuroimmune interactions. Future work should examine whether direct manipulation of glial activity in the piriform cortex can rescue olfactory deficits or normalize behavioral responses to aversive stimuli. Such interventions may yield novel therapeutic strategies that target glial function to alleviate sensory and affective symptoms in depression.

## Acknowledgements

We thank Angles Salles and Janet Richmond for sharing equipment and resources. The Department of Biological Sciences, Imaging Core for microscopy assistance, and all members of the Zak Laboratory for helpful discussions throughout this project.

## Extended Data

**Figure 6-1.**
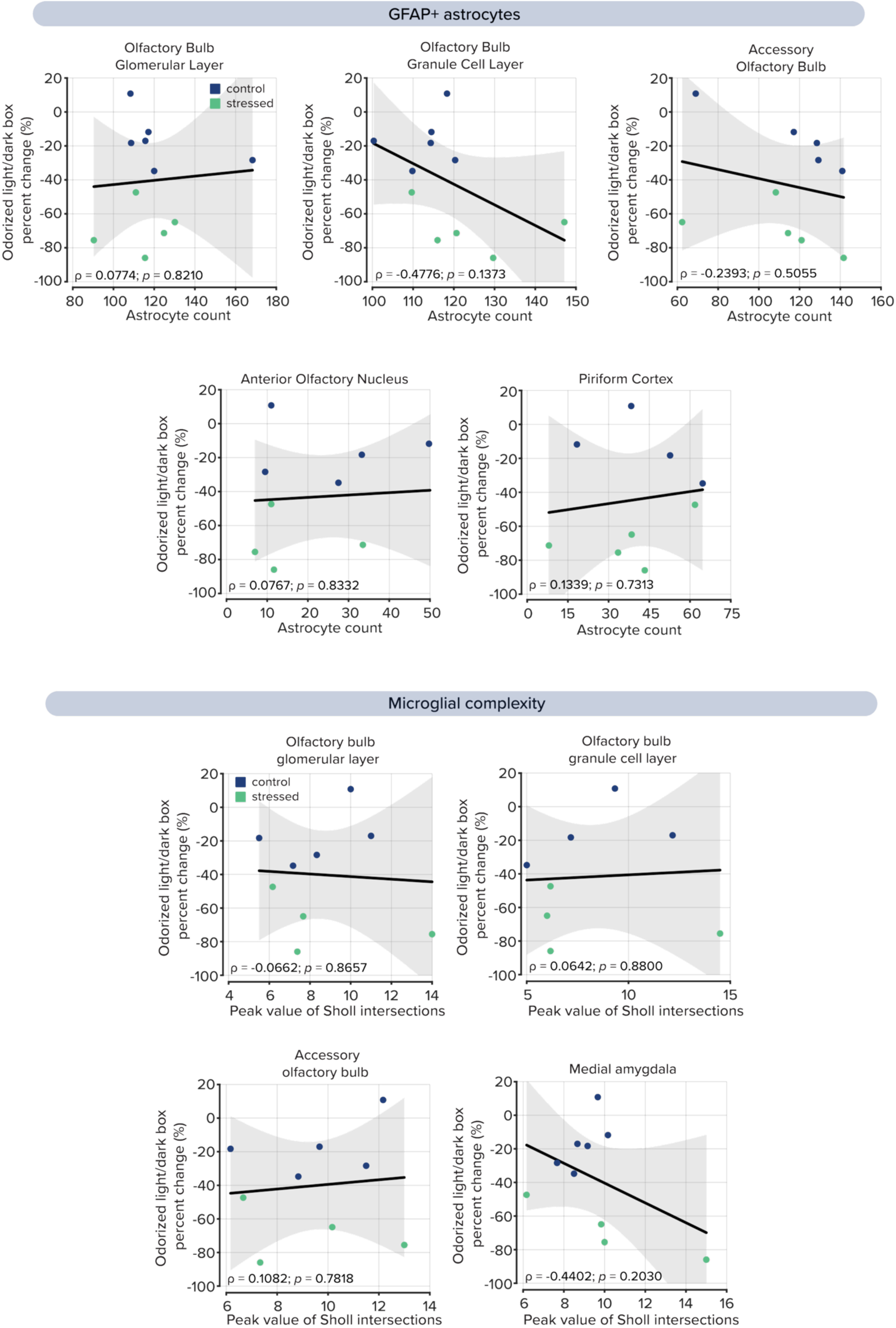
Correlation analysis between labeled astrocytes and odorant aversion in the olfactory bulb glomerular layer, olfactory bulb granule cell layer, accessory olfactory bulb, anterior olfactory nucleus, and anterior piriform cortex.

**Figure 6-2.**
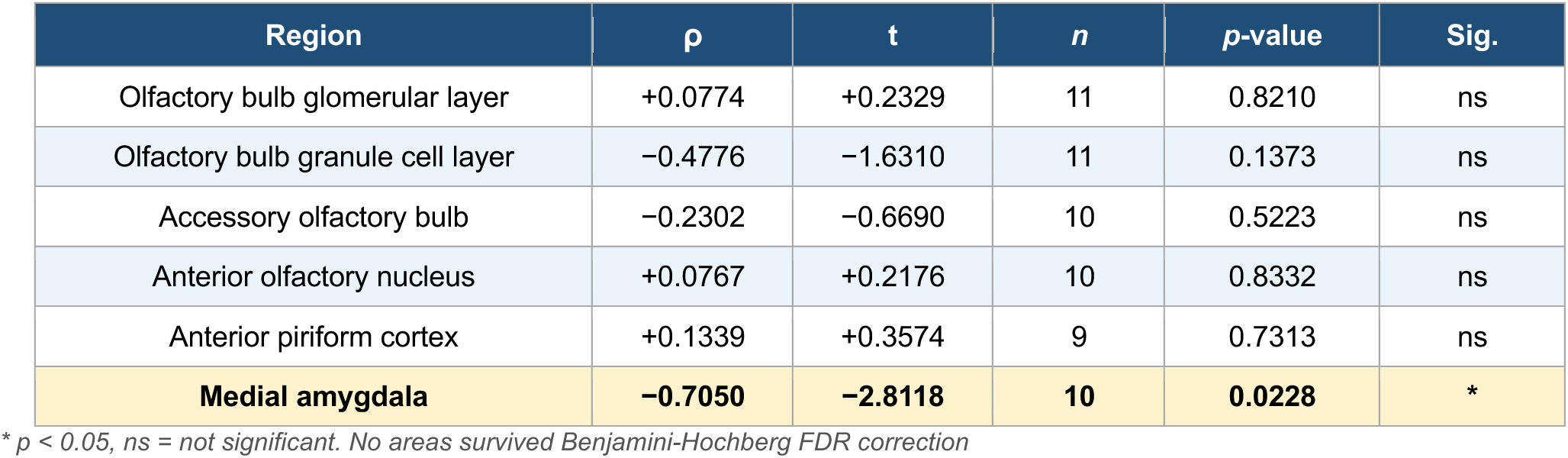
Pearson correlations between odor avoidance behavior and astrocyte counts per animal, pooled across groups.

**Figure 6-3.**
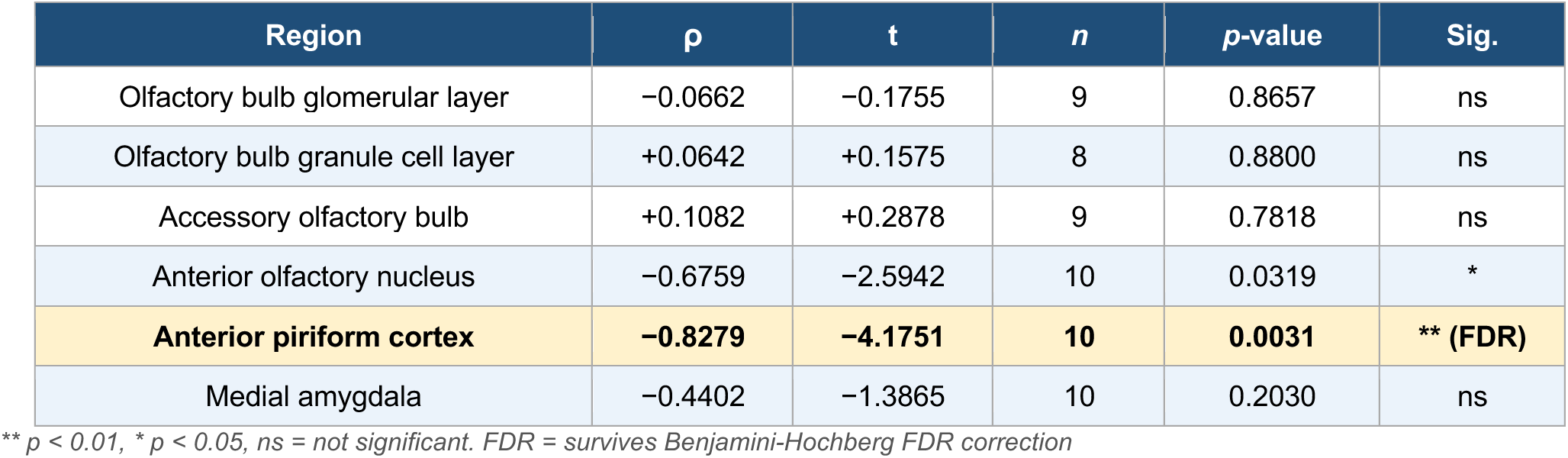
Pearson correlations between odor avoidance behavior and peak Sholl intersections per animal, pooled across groups

